# Sex differences in the genetic architecture of obsessive-compulsive disorder

**DOI:** 10.1101/219170

**Authors:** Ekaterina A. Khramtsova, Raphael Heldman, Eske M. Derks, Dongmei Yu, TS/OCD Psychiatric Genomics Disorders Workgroup, Lea K. Davis, Barbara E. Stranger

**Affiliations:** Section of Genetic Medicine, Department of Medicine, University of Chicago, Chicago, IL; Institute for Genomics and Systems Biology, University of Chicago, Chicago, IL; University of California, Berkeley, California; Queensland Institute of Medical Research, Brisbane, Australia; Psychiatric and Neurodevelopmental Genetics Unit, Center for Genomic Medicine, Department of Psychiatry, Harvard Medical School, Massachusetts General Hospital, Boston, Massachusetts; Vanderbilt Genetics Institute; Vanderbilt University Medical Center, Nashville, TN; Division of Medical Genetics, Department of Medicine, Vanderbilt University Medical Center, Nashville, TN; Department of Psychiatry and Behavioral Sciences, Vanderbilt University Medical Center, Nashville, TN; Center for Data Intensive Science, University of Chicago, Chicago, IL

## Abstract

Obsessive-compulsive disorder (OCD), a highly heritable complex phenotype, demonstrates sexual dimorphism in age of onset and clinical presentation, suggesting a possible sex difference in underlying genetic architecture. We present the first genome-wide characterization of the sex-specific genetic architecture of OCD, utilizing the largest set of OCD cases and controls available from the Psychiatric Genomics Consortium. We assessed evidence for several mechanisms that may contribute to sexual-dimorphism including a sexually dimorphic liability threshold, the presence of individual sex-specific risk variants on the autosomes and the X chromosome, genetic and phenotypic heterogeneity, and sex-specific pleiotropic effects. We observed a strong genetic correlation between male and female OCD and no evidence for a sexually dimorphic liability threshold model. While we did not detect any sex-specific genome-wide associations, we observed that the SNPs with sexually dimorphic effects showed an enrichment of regulatory variants influencing expression of genes in immune tissues. Furthermore, top sex-specific genome-wide associations were enriched for regulatory variants in different tissues, suggesting evidence for potential sex difference in the biology underlying risk for OCD. These findings suggest that future studies with larger sample sizes hold great promise for the identification of sex-specific risk factors for OCD, significantly advancing our understanding of the differences in the genetic basis of sexually dimorphic neuropsychiatric traits.

## Introduction

Many human diseases exhibit sexual dimorphism in onset, prevalence, prognosis, and clinical features, however, the etiology of these differences remains poorly understood. Studies have demonstrated a contribution to sex-biased phenotypes from autosomal genetic variation (Ober, Loisel, and Gilad 2008; Mackay 2004; Korstanje et al. 2004), motivating characterization of the sex-specific genetic architecture of complex traits. Neuropsychiatric disorders such as bipolar disorder, autism spectrum disorder, attention deficit hyperactivity disorder, schizophrenia, Tourette syndrome (TS), obsessive-compulsive disorder (OCD), and depression, display sex-bias in age of onset, progression, and/or prevalence. The genetic basis of sexual dimorphism in OCD has not yet been explored.

The SNP-based heritability of OCD (Pauls et al. 2014) is approximately 24-32% (International Obsessive Compulsive Disorder Foundation Genetics Collaborative (IOCDF-GC) and OCD Collaborative Genetics Association Studies (OCGAS) 2017). OCD also demonstrates earlier onset in males (Flament et al. 1990; Swedo et al. 1989; Bellodi et al. 1992; Boileau 2011), and sex-biased clinical symptom presentation. Epidemiological studies indicate a worldwide lifetime prevalence of OCD between 1 and 3% (Kessler et al. 2005; Ruscio et al. 2010; Torres and Lima 2005; Weissman et al. 1994) and while boys comprise approximately two thirds of the childhood cases of OCD, typically defined as onset before age 15 (Flament et al. 1990; Swedo et al. 1989; Bellodi et al. 1992; Boileau 2011), the lifetime prevalence of OCD measured in adults is equivalent between the sexes. Females with OCD, in addition to demonstrating a later age of onset also have lower familial loading and higher rates of precipitating events, including pregnancy and childbirth (Millet et al. 2004). Compared to females, males with OCD report more religious, sexual, and symmetry symptoms, more alcohol dependence, and lower rates of marriage and employment. Females with OCD are more likely to be married, report more sexual abuse during childhood, often report exacerbation of symptoms in the premenstrual/menstrual period, during/shortly after pregnancy, with menopause, and tend to have more contamination and cleaning compulsions, as well as eating disorders, reviewed in Mathis et al (Mathis et al. 2011).

Although the recently-published genome-wide association studies of OCD (Mattheisen et al. 2015; Stewart et al. 2013; International Obsessive Compulsive Disorder Foundation Genetics Collaborative (IOCDF-GC) and OCD Collaborative Genetics Association Studies (OCGAS) 2017) have not discovered genome-wide significant associations, these studies have demonstrated that common variants account for a significant proportion of OCD heritability (Davis et al. 2013). They also indicate that the strongest associated variants in OCD GWAS are enriched for expression quantitative trait loci (eQTLs) and methylation QTLs derived from frontal lobe, cerebellum, and parietal lobe tissue (Stewart et al. 2013), demonstrating that biologically meaningful associations exist within the top ranked SNPs and that increasing sample sizes will likely identify significant common variant associations for OCD risk. In addition to increasing sample size, another approach to improve power for GWAS is reducing heterogeneity. Given the difference in clinical presentation of OCD in males and females, we wished to test the hypothesis that the genetic architecture varies between the sexes. If true, this could provide an additional approach for improving power for OCD gene-finding efforts.

In this study, we sought to characterize the sex-specific genetic architecture of OCD using multiple approaches. Several studies have demonstrated the value of performing sex-stratified GWAS, assessing heterogeneity of effects between sexes, or including a genotype-sex interaction term in routine GWAS, as these approaches have discovered novel loci which were previously undetected due to heterogeneity between sexes (Mitra et al. 2016; Martin et al. 2017; Taylor et al. 2013; Randall et al. 2013). Thus, we first performed a sex-stratified genome-wide association meta-analysis and genotype-sex interaction meta-analysis including autosomes and the X chromosome. We then developed an approach to identify SNPs with Sexually Dimorphic Effect (SDEs), and assessed whether the SDEs regulate gene expression and are enriched for associations with sexually dimorphic anthropometric traits (i.e. height, weight, body mass index, hip and waist circumference) as observed in autism spectrum disorders (Mitra et al. 2016). Third, we performed SNP-based heritability analysis to (a) assess the proportion of overall OCD heritability explained by the X chromosome, and (b) test for evidence of variable liability threshold for OCD between males and females. This phenomenon, known as the Carter effect, in which the sex with the lower prevalence/milder presentation requires a higher genetic burden to become affected, has been reported for several complex traits (Kruse et al. 2012). Fourth, we performed a sex-stratified genetic correlation analysis with other traits which may play a role in OCD development (e.g. brain volumes), show sexual dimorphism (e.g. autism, Tourette syndrome, attention deficit hyperactivity disorder, etc.), or are known to show differences in comorbidity between males and females with OCD (e.g. smoking, eating disorders, and reproductive behavior). Here, we present the first genome-wide assessment of the sex-specific genetic architecture of OCD utilizing the largest OCD dataset currently available. We also provide best practices for sex-stratified analysis which can be adopted in future studies of OCD and other phenotypes.

## Methods

### Datasets and software

The datasets (Supplementary Figure 1) used in this study comprise the OCD Psychiatric Genomics Consortium sample and are fully described in primary publications (Stewart et al. 2013; Mattheisen et al. 2015; International Obsessive Compulsive Disorder Foundation Genetics Collaborative (IOCDF-GC) and OCD Collaborative Genetics Association Studies (OCGAS) 2017). All participants over 18 and the parents of participants under 18 gave written informed consent and this work was approved by the relevant institutional review boards at all participating sites. Participants of European ancestry were selected for this study and include cases and controls from Dutch, South African, European, and Ashkenazi Jewish ancestries. Additionally, trio samples were included in the meta-analysis and consisted of proband cases and pseudo-controls. The pseudo-controls were derived from the parental haplotypes that were not transmitted from parents to probands.

### Sample and genotype level quality control and imputation

#### Autosomes

Genotype level data from all studies were pre-phased with SHAPEIT2 (Delaneau, Zagury, and Marchini 2013), and imputed to the 1000 Genomes Project reference panel (Phase I integrated variant set release; NCBI build 37 (hg19)) using IMPUTE2 (Howie, Marchini, and Stephens 2011), using the Ricopili pipeline (Schizophrenia Working Group of the Psychiatric Genomics Consortium 2014). Prior to imputation, SNPs with call rate<0.98, minor allele frequency (MAF)<0.01, case-control differential missingness>0.02, Hardy–Weinberg equilibrium (HWE) p-values <1e-6 for controls and <1e-10 for cases were removed using PLINK (Purcell et al. 2007). After imputation, any SNPs with IMPUTE2 info score <0.6 and certainty <0.8 were removed. After splitting the data sets by sex, SNPs with MAF <0.05 were removed from each sex.

At the individual level, samples were removed if they had a call rate <0.98, the absolute value of heterozygosity (F_HET)>0.20, or showed an inconsistency between genetic sex and reported sex. Furthermore, pairwise identity by descent (IBD) analysis was used to identify cryptically related individuals, and one individual was removed at random from any pair related at the approximate level of first cousins (pi-hat>0.2). Principal component analyses were performed separately for each sub-population, (Supplementary Methods; Supplementary Figure 2 and 3) and case/control matching was performed separately for each sex using EIGENSOFT (Price et al. 2006). After quality control, the total sample comprised 4,038 males and 5,832 females. The numbers of post-QC SNPs and individuals are listed in Supplementary Table 1.

**Table 1.**
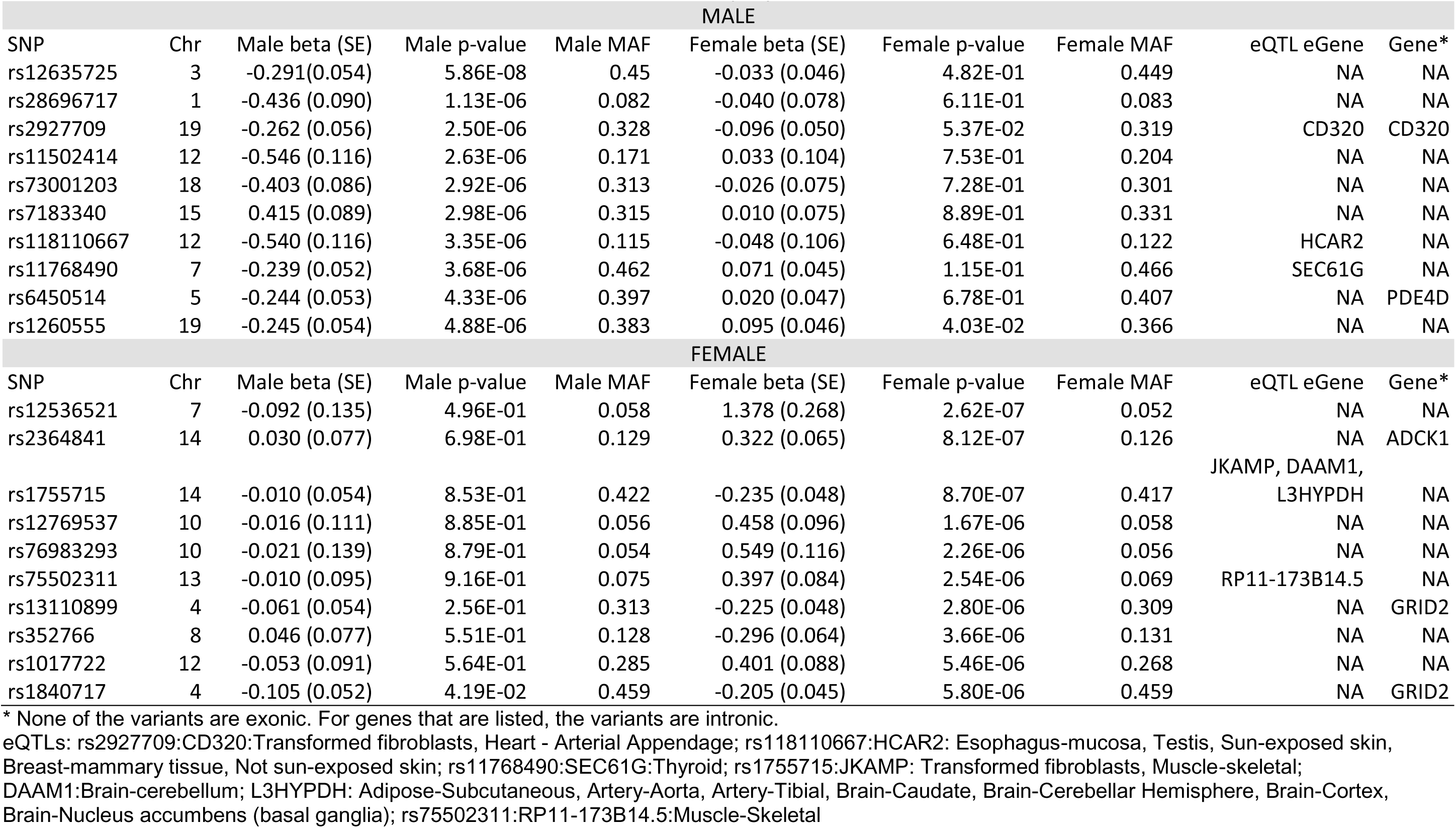
Top ten LD-independent (r_2_=0.2) associations in male-specific and female-specific genome-wide association studies. For each variant, both female and male association betas and p-values are shown. All variants are annotated as intergenic; however, none are exonic. Each variant that is an eQTL is labeled with the target gene(s), with the source tissue listed in the table footnote. Abbreviations: MAF, minor allele frequency; Chr, chromosome; SE, standard error; eQTL, expression quantitative trait locus; eGene, eQTL target gene.

#### X chromosome

X chromosome genotypes were processed separately from autosomal genotypes as additional care is required for pre-phasing, imputation, and post-imputation QC. At the genotype level, the pre-imputation QC steps for the X chromosome SNPs were the same as for the autosomes. An additional flag of ‐chrX was added when running SHAPEIT2 and IMPUTE2 software. Post-imputation, we employed the XWAS QC pipeline to remove variants in the pseudoautosomal regions (PARs), variants that were not in Hardy-Weinberg equilibrium in females, or variants with significantly different MAF (p<0.05/#SNPs) and differential missingness (P<10^−7^) between males and female controls (Gao et al. 2015).

For imputation, we included those samples that passed both autosomal QC, and had a call rate >0.98 on the X chromosome. Furthermore, because we could not use the same case/pseudo-control design for the trio data (i.e. due to lack of a non-transmitted X chromosome from the fathers of affected females), we included only the affected individuals from the trio data, ancestry-matched them to controls, and analyzed them with the case/control data. We performed PCA and removed any trio cases without matching controls.

### Genome-wide association meta-analysis

For each individual dataset, we performed sex-stratified GWAS. Principal components demonstrating association with the phenotype were included as covariates. We then used the inverse variance method implemented in METAL (Willer, Li, and Abecasis 2010) to meta-analyze summary statistics of each dataset for sex-stratified analysis. Furthermore, we performed GWAS and meta-analysis on the combined male/female sample for each subpopulation to ensure that our sex-specific QC yielded results consistent with the recently reported OCD meta-GWAS using the same data (International Obsessive Compulsive Disorder Foundation Genetics Collaborative (IOCDF-GC) and OCD Collaborative Genetics Association Studies (OCGAS) 2017). The correlation calculated using LD score regression (B. K. Bulik-Sullivan et al. 2015) between our meta-analysis and the previously published meta-analysis (International Obsessive Compulsive Disorder Foundation Genetics Collaborative (IOCDF-GC) and OCD Collaborative Genetics Association Studies (OCGAS) 2017) was 1.0521, se=0.0141. A genetic correlation greater than one can be observed when there is sample/reference LD Score mismatch or model misspecification (e.g., low LD variants have slightly higher h^2^per SNP).

### Genotype-sex interaction analysis

We used PLINK to perform a genotype-sex (GxS) interaction analysis, with principal components as covariates, in each of the individual datasets. We then used METAL to meta-analyze the interaction results.

### Assessment of heterogeneity from sex-stratified GWAS

We used Z-scores (correlated with Cochran's Q statistic but provides directionality of the effect, Supplementary Methods) to assess heterogeneity between males and females. To obtain a Z-score for each tested variant, we calculated the differences in effect sizes (beta) between the sexes weighted by the square root of the sum of beta standard errors squared (equation 1).

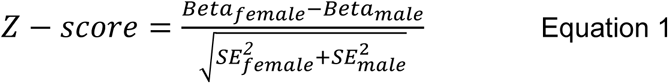

We define SNPs with Sexually Dimorphic Effect (SDEs) as those variants at the extreme ends of the distribution with an absolute value of the Z-score greater than 3 (|Z-score|>3), which is roughly equivalent to p<10^−3^, and represents 0.3% of all tested SNPs

### Heritability estimates and genetic correlation

To calculate the sex-specific narrow-sense SNP-based heritability (h^2^), (i.e., the proportion of phenotypic variation attributable to the additive effect of all SNP variants in each sex), we used two methods: 1) LD score regression (LDSC) as implemented in LDSC v1.0.0 (B. K. Bulik-Sullivan et al. 2015) and 2) restricted maximum likelihood analysis (REML) implemented in GCTA v1.24.4 (Yang et al. 2011). LDSC analysis was performed on the sex-stratified meta-analysis summary statistics from all study datasets. Meta-analyzed imputed SNPs which overlapped with a panel of high confidence HapMap SNPs were used for the LD score regression. Because our data set is composed of European individuals, we downloaded precomputed LD scores (B. K. Bulik-Sullivan et al. 2015; B. Bulik-Sullivan et al. 2015). Using all individuals, we calculated the total and sex-stratified heritability, checked for residual population stratification (based on the LDSC intercept (B. K. Bulik-Sullivan et al. 2015)), and calculated the genetic correlation between males and females. A range of 1-3% OCD population prevalence was used to transform from the observed heritability scale to the liability scale.

For REML analysis, we used a combination of the IOCDF-GC and OCGAS European data sets plus the cases from the IOCDF-GC and OCGAS trio data set and performed an additional PCA analysis on this combined sample to remove any outliers. Genetic relationship matrices (GRM) for autosomes and chromosome X were generated for combined and sex-stratified datasets, removing any individuals who are closely related (IBD>0.05). All pruned imputed SNPs were used to determine the top 20 principal components using smartpca in EIGENSOFT (Price et al. 2006). Genomic-relatedness-based restricted maximum-likelihood (GREML) analysis was performed on the autosomes and the X chromosome (taking into account dosage compensation, Supplementary Methods) using GRMs and the top 20 ancestry covariates to estimate the proportion of phenotypic variation accounted for by the variants. Because the prevalence of OCD reported in the literature ranges from 1-3%, we estimated heritability on the liability scale using OCD prevalence of 1%, 1.5%, 2%, and 3%. Bivariate GREML analysis was performed in GCTA to assess genetic correlation between sexes. To assess the proportion of the total heritability contributed by the X chromosome, a separate GRM was generated for each of the 23 chromosomes. Then, all chromosomes were analyzed jointly in a single GREML analysis with 20 PCs to account for population substructure.

### Enrichment of expression quantitative trait loci in brain and immune tissues among OCD-associated variants and SDEs and their implication in biological processes

To assess eQTL enrichment, specifically to test for an enrichment for a gene regulatory role among top GWAS associations and SDEs, we quantified the enrichment of the number of eQTL target genes (eGenes) associated with OCD-associated SNPs. Expression quantitative trait loci (eQTL) enrichment analysis was performed on (a) SNPs nominally associated with OCD (p<10−3) in the combined and sex-stratified GWAS analysis, and (b) SDEs. Prior to clumping (r^2^=0.2, 500kb window), each set of SNPs was also filtered for variants with fewer than five hundred individuals present in the meta-analysis. We also report results of analyses of unfiltered SNPs (Supplementary Figure 7). eQTL annotation was performed using previously published eQTL results (Supplementary Table 2), including eQTLs derived from 10 regions of the brain and whole blood from GTEx (GTEx Consortium et al. 2017), a meta-eQTL analysis of brain cortex tissue (Kim et al. 2014), as well as CD4+ T cells and CD14+ monocytes (Raj et al. 2014). To assess eQTL enrichment, 1000 randomly ascertained SNPs sets were generated using SNPsnap (Pers, Timshel, and Hirschhorn 2015), sampled without replacement from the European catalogue of 1000 Genomes SNPs, and matched for minor allele frequency (± 5%), gene density (± 50%), distance to nearest gene (within a 1000kb window), and LD buddies (± 50%) at r^2^=0.8.

SNPs in the OCD-associated set and the null matched SNP sets were annotated both with *cis*-eQTL status and with the genes they regulate (i.e., eGenes) in various brain and immune tissues. The enrichment p-value was calculated as the proportion of randomized sets in which the number of eGenes matched or exceeded the observed count among trait-associated SNPs. If multiple variants implicated the same eGene in a tissue or cell type, the eGene was counted only once. This strategy is different from counting individual eQTLs variants, as was done for the previous OCD GWAS (Stewart et al. 2013), where multiple SNPs may be regulating the same gene and thus over-counted, while here all eQTLs targeting the same gene are counted only once. We also performed “pan-tissue” eQTL eGene analysis by combining the eQTL results from all the brain tissue subtypes and all the immune tissue and cell subtypes. If an eGene was present in more than one tissue, it was counted only once. To exclude the possibility of eQTL enrichment overestimation due to the gene-rich MHC region, we performed eQTL enrichment analysis both including and excluding SNPs in the HLA region. The enrichment was considered significant if the empirical p-value exceeded Bonferroni multiple testing correction threshold p<0.0036 (i.e. 0.05/14 tissues).

To evaluate underlying biological pathways that may be regulated by the eQTLs identified in the brain and immune tissues, we applied Gene Ontology (GO) enrichment analyses to the eGene sets as implemented in the Gene Set Enrichment Analysis (GSEA, Broad Institute) (Mootha et al. 2003; Subramanian et al. 2005).

### Enrichment of OCD-associated SNPs among anthropometric trait SDEs

We tested for enrichment of anthropometric traits SDEs (ASDEs) among SNPs nominally associated with OCD (p<10^−3^) in (a) the combined male/female analysis, (b) the sex-stratified analyses, and (c) the SDEs. ASDEs were defined using the approach described in (Mitra et al. 2016) (Z-score p<=10^−3^) for several anthropomorphic traits from the GIANT consortium (Randall et al. 2013): weight, height, body mass index (BMI), hip circumference (HIP), HIP adjusted for BMI (HIPadjBMI), waist circumference (WC), WC adjusted for BMI (WCadjBMI), waist-to-hip ratio (WHR), and WHR adjusted for BMI (WHRadjBMI) resulting in a total of 12,006 unique ASDEs identified across GIANT phenotypes. We determined the overlap of ASDEs with each OCD subset (Supplementary Figure 8), as well as with 1000 matching SNP sets for each of the OCD subsets. An empirical enrichment p-value was calculated as the proportion of null randomized sets in which the overlap matched or exceeded the observed overlap using the OCD associated SNPs.

### Sex-stratified genetic correlation analyses

Genetic correlation analysis of OCD with several phenotypes of interest was performed for the combined sample and sex-stratified samples using LD score regression (B. K. Bulik-Sullivan et al. 2015). Summary statistics for the following phenotypes were obtained: Tourette Syndrome (Scharf et al. 2013; Yu et al. 2015), obsessive-compulsive symptoms (den Braber et al. 2016), attention deficit hyperactivity disorder (Neale et al. 2010), autism (unpublished, available via Psychiatric Genomics Consortium), bipolar disorder (Psychiatric GWAS Consortium Bipolar Disorder Working Group 2011), major depressive disorder (Major Depressive Disorder Working Group of the Psychiatric GWAS Consortium et al. 2013), schizophrenia (Schizophrenia Working Group of the Psychiatric Genomics Consortium 2014), anxiety disorders (Otowa et al. 2016), depressive symptoms, neuroticism, subjective well-being (Okbay et al. 2016), anorexia (Boraska et al. 2014), body mass index (Speliotes et al. 2010), tobacco usage (Tobacco and Genetics Consortium 2010), reproductive behavior (Barban et al. 2016) and structural brain measures (accumbens, amygdala, pallidum, caudate, thalamus, putamen volumes) (Hibar et al. 2015), hippocampal volume (Hibar et al. 2017), and intracranial volume (Adams et al. 2016). We identified high confidence HapMap SNPs (for which the LD scores have been precomputed) present in the OCD summary statistics and each of the other summary statistics. For continuous traits (e.g. cognitive performance, brain structure volumes) no sample or population prevalence was specified. For binary traits the sample prevalence was calculated based on the reported number of cases in the sample, while the population prevalence was obtained from the literature (Supplementary Table 7).

## Results

### Sex-stratified genome-wide association and sex by genotype interaction analyses

Genomic control lambda (λ_GC_) revealed no significant evidence of population stratification in the male-specific (λ_GC_=1.019), the female-specific (λ_GC_=1.026), or the combined (λ_GC_=1.051) meta-analyses. The intercepts estimated by LD score regression of 1.0016, 0.9907, 1.0048 for sex-combined, female-only, and male-only, respectively, suggested that the mild inflation observed on the quantile-quantile plots (λ_GC_) was not due to population stratification but rather to polygenic effects. The Manhattan and quantile-quantile plots (Figure 1 A-B) demonstrated no genome-wide significant associations in either males or females. Furthermore, a scatter plot of - log10(p-values) for sex-specific genome-wide associations indicated little overlap in the top signals across sexes (Figure 1C, Table 1).

**Figure 1.**
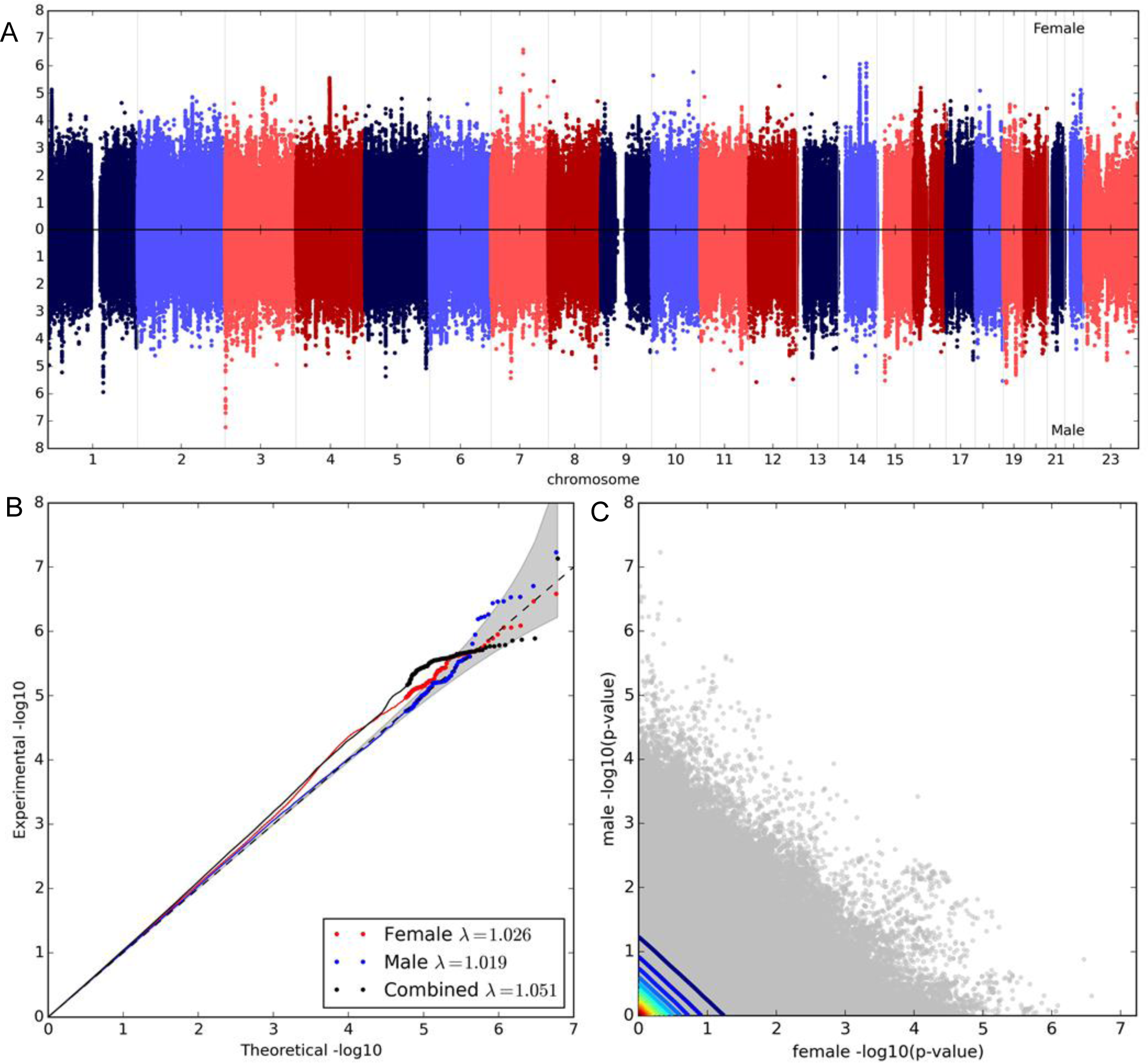
Manhattan and quantile-quantile plots for sex-stratified meta-GWAS. Meta-GWAS was run separately for females (1525 cases and 4307 controls) and males (1249 cases and 2789 controls) on ~5.5 million imputed SNPs (MAF>5%). (A) The peaks pointing up on the plot are the results for female analysis and the peaks pointing down are the results for male analysis. Although not genome-wide significant, several suggestive peaks can be observed in one sex and not observed in the other. (B) Quantile-quantile plot for sex-stratified and combined meta-GWAS. (C) −log(p-value) for SNP association in females plotted against males. Contour lines colored from red to blue indicate decreasing data density.

We calculated a Z-score and its p-value for each SNP to assess heterogeneity in effect size between male‐ and female-specific associations (top ten SDEs, Table 2). The QQ plots of Z-score p-values indicated no enrichment for SDEs (Supplementary Figure 4), and that the difference in effect size for SDEs was not driven by minor allele frequency (MAF) differences between sexes (Supplementary Figure 5). The MAF distributions for SDEs and all tested SNPs were identical, and sexually-dimorphic loci were distributed across the genome proportional to chromosome length (Supplementary Figure 5D). P-values from a sex by genotype interaction test were highly correlated with Z-score p-values from the sex-stratified analysis (autosomal SNPs Pearson's r=0.65, p<2.2e-16, X chromosome SNPs Pearson's r=0.71, p<2.2e-16). Furthermore, GWAS results in the combined sample with or without sex as a covariate were highly correlated (LDSC r_g_=0.9994, se=0.0005).

**Table 2.** Top ten LD-independent (r^2^=0.2) SDEs. For each SDE, the female and male association betas and p-values, z-score and its p-value, and the genotype-sex interaction p-value are shown. All Variants are annotated as intergenic; however, none are exonic. Each variant that is an eQTL is labeled with the target gene(s), with the source tissue listed in the table footnote. Abbreviations: SDEs, SNPs with Sexually Dimorphic Effect; MAF, minor allele frequency; Chr, chromosome; SE, standard error; eQTL, expression quantitative trait locus; eGene, eQTL target gene.

| SNP | Chr | Male<br>beta (SE) | Male<br>p-value | Male<br>MAF | Female<br>beta (SE) | Female<br>p-value | Female<br>MAF | eQTL<br>eGene | SDE z-score | SDE z-score<br>p-value | GxS interaction<br>p-value | Gene* |
| --- | --- | --- | --- | --- | --- | --- | --- | --- | --- | --- | --- | --- |
| rs12536521 | 7 | -0.09 (0.14) | 4.96E-01 | 0.058 | 1.38 (0.27) | 2.62E-07 | 0.052 | NA | 4.905 | 9.33E-07 | 1.60E-01 | NA |
| rs4798525 | 18 | 0.28 (0.06) | 2.13E-05 | 0.224 | -0.14 (0.06) | 1.27E-02 | 0.228 | NA | -4.849 | 1.24E-06 | 2.06E-02 | LAMA1 |
| rs2077613 | 17 | 0.22 (0.05) | 1.72E-05 | 0.425 | -0.11 (0.05) | 1.82E-02 | 0.427 | NA | -4.795 | 1.63E-06 | 4.36E-06 | NA |
| rs11064706 | 12 | -0.21 (0.07) | 4.44E-03 | 0.157 | 0.25 (0.06) | 5.97E-05 | 0.153 | NA | 4.762 | 1.92E-06 | 1.94E-01 | NA |
| rs79886445 | 6 | -0.30 (0.10) | 2.57E-03 | 0.084 | 0.30 (0.08) | 1.88E-04 | 0.081 | NA | 4.694 | 2.68E-06 | 4.16E-06 | NA |
| rs17815599 | 3 | -0.27 (0.06) | 2.45E-05 | 0.211 | 0.13 (0.06) | 2.44E-02 | 0.209 | NA | 4.64 | 3.48E-06 | 3.40E-05 | NA |
| rs11119584 | 1 | 0.18 (0.05) | 8.54E-04 | 0.408 | -0.15 (0.05) | 1.46E-03 | 0.391 | NA | -4.603 | 4.16E-06 | 3.35E-04 | IL19 |
| rs6034007 | 20 | -0.13 (0.05) | 1.08E-02 | 0.438 | 0.18 (0.05) | 5.95E-05 | 0.435 | NA | 4.561 | 5.08E-06 | 1.19E-05 | MACROD2 |
| rs7334430 | 13 | -0.29 (0.08) | 5.47E-04 | 0.124 | 0.21 (0.07) | 3.08E-03 | 0.122 | NA | 4.549 | 5.38E-06 | 1.41E-05 | NA |
| rs11768490 | 7 | -0.24 (0.05) | 3.68E-06 | 0.462 | 0.07 (0.05) | 1.15E-01 | 0.466 | SEC61G | 4.526 | 6.02E-06 | 3.78E-04 | NA |

\* None of the variants are exonic. For genes that are listed, the variants are intronic. eQTLs: rs11768490:SEC61G:Thyroid

### Genetic correlation for OCD is high between males and females

For highly polygenic traits, individual genetic variants, including the most significantly associated variants, typically explain only a small fraction of a trait's phenotypic variance. To characterize the sex-specific genetic architecture of OCD, we explored sub-threshold associations and their contribution to OCD heritability (h^2^).

The difference in heritability estimates (Table 3) between males (
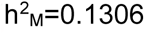
, SE = 0.0966) and females (
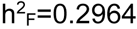
, SE = 0.0787), as determined by LDSC regression, was not statistically significant, and the genetic correlation between the sexes was substantial (r_g_ = 1.0427, SE = 0.5089, p=0.0405). The restricted maximum likelihood analysis (REML) estimates of heritability were almost identical between males (
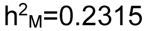
, se=0.0717, p=0.0011) and females (
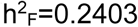
, se=0.0569, p=1.07e-05), and to the combined estimate (h^2^=0.2376, se=0.0333, p=8.621e-14). The REML genetic correlation between males and females was 1.00 (se=0.27). The observed patterns were also robust across population prevalence rates (Supplementary Table 3).

**Table 3.** Sex-stratified and combined heritability estimates for OCD from autosomes and the X chromosome. Abbreviations: OCD, obsessive-compulsive disorder; LDSC, linkage disequilibrium score regression; GCTA, genome-wide complex trait analysis; N, number of individuals in the analysis; h^2^, SNP-heritability; SE, standard error.

| Condition | LDSC |  |  | GCTA |  |  |  |  |  |  |  |
| --- | --- | --- | --- | --- | --- | --- | --- | --- | --- | --- | --- |
|  | Autosomes |  |  | Autosomes |  |  |  | X Chromosome |  |  |  |
| | N | $h^2$ | SE | N | $h^2$ | SE | P-value | N | $h^2$ | SE | P-value |
| Combined | 9,870 | 0.2246 | 0.0454 | 7,051 | 0.2376 | 0.0333 | 8.62E-14 | 7,059 | 0.0104 | 0.0049 | 6.20E-03 |
| Male | 4,038 | 0.1306 | 0.0966 | 2,781 | 0.2315 | 0.0761 | 1.10E-03 | 2,778 | 0.028 | 0.0129 | 9.80E-03 |
| Female | 5,832 | 0.2964 | 0.0787 | 4,274 | 0.2403 | 0.0569 | 1.07E-05 | 4,281 | 0.0141 | 0.0083 | 2.72E-02 |

### X chromosome contributes to the polygenic architecture of OCD in both sexes

One of the mechanisms by which sex differences in OCD could arise is through genetic risk deriving from the sex chromosomes. We observed no significant associations on the X chromosome in either the combined or sex-stratified analyses. A QQ-plot indicated that there was no excess of SDEs on the X chromosome (Supplementary Figure 4). Using REML, we estimated the X chromosome (1.6% of total SNPs) heritability as 
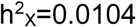
 (se=0.0049, p=0.006244), which comprised 3.8% of total OCD heritability, and was consistent with expectation (Supplementary Figure 6). When analyzed in each sex separately, X chromosome heritability was not statistically different between females (
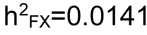
, se=0.0083, p=0.0271) and males (
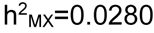
, se=0.0128, p=0.0098) at 2.5% OCD prevalence. Results were again robust to estimates derived using a range of OCD prevalence (Supplementary Table 3).

### eQTL enrichment observed among SDEs and strongest associations from sex-stratified GWAS

To investigate the functional effects of top associations (p<10^−3^) from the sex-stratified GWAS analysis and SDEs, we annotated each SNP as to whether it was an expression quantitative trait locus (eQTL) using publicly-available brain and immune eQTLs. Specifically, we tested for an enrichment for a gene regulatory role for OCD-associated SNPs, as quantified by an enrichment of the number of eQTL target genes (eGenes) associated with OCD-associated SNPs compared to random, matched SNPs. We tested for enrichment of eQTLs derived from brain tissues because brain is the primary tissue of interest, but also eQTLs derived from immune cells because the immune system has been previously implicated in several neuropsychiatric and neurodegenerative traits (Schizophrenia Working Group of the Psychiatric Genomics Consortium 2014; Marsh et al. 2016; Heneka, Golenbock, and Latz 2015; Furtado and Katzman 2015a), including OCD (Furtado and Katzman 2015b; Murphy, Sajid, and Goodman 2006) For eQTL annotation, we used previously published eQTLs datasets derived from 10 brain regions and whole blood from GTEx (GTEx Consortium et al. 2017), as well as CD4+ T cells and CD14+ monocytes (Raj et al. 2014), as a proxy for adaptive and innate immune responses, respectively.

SDEs showed a significant enrichment for eQTLs from CD4+ T cells (p=0.001), whole blood (p<0.001), and the combination of immune tissues (p<0.001) (Figure 2, Supplementary Table 4). 122 eGenes were implicated by brain eQTLs and 220 by immune eQTLs, with 31 eGenes deriving from both tissues (Supplementary Table 5). Top female associations were enriched for brain eQTLs (p=0.003 when excluding the cerebellum, but p=0.009 when including the functionally distinct cerebellum). Top male associations were enriched for immune eQTLs (p=0.003). The most significant associations from the combined male/female GWAS did not show an eQTL enrichment in any of the tissues examined. Including HLA SNPs resulted in a weaker enrichment for the top male (p_IMMUNE_=0.015) and female (p_BRAIN_=0.008) associations, but did not affect the combined male/female GWAS or SDEs eQTL enrichment analyses.

**Figure 2.**
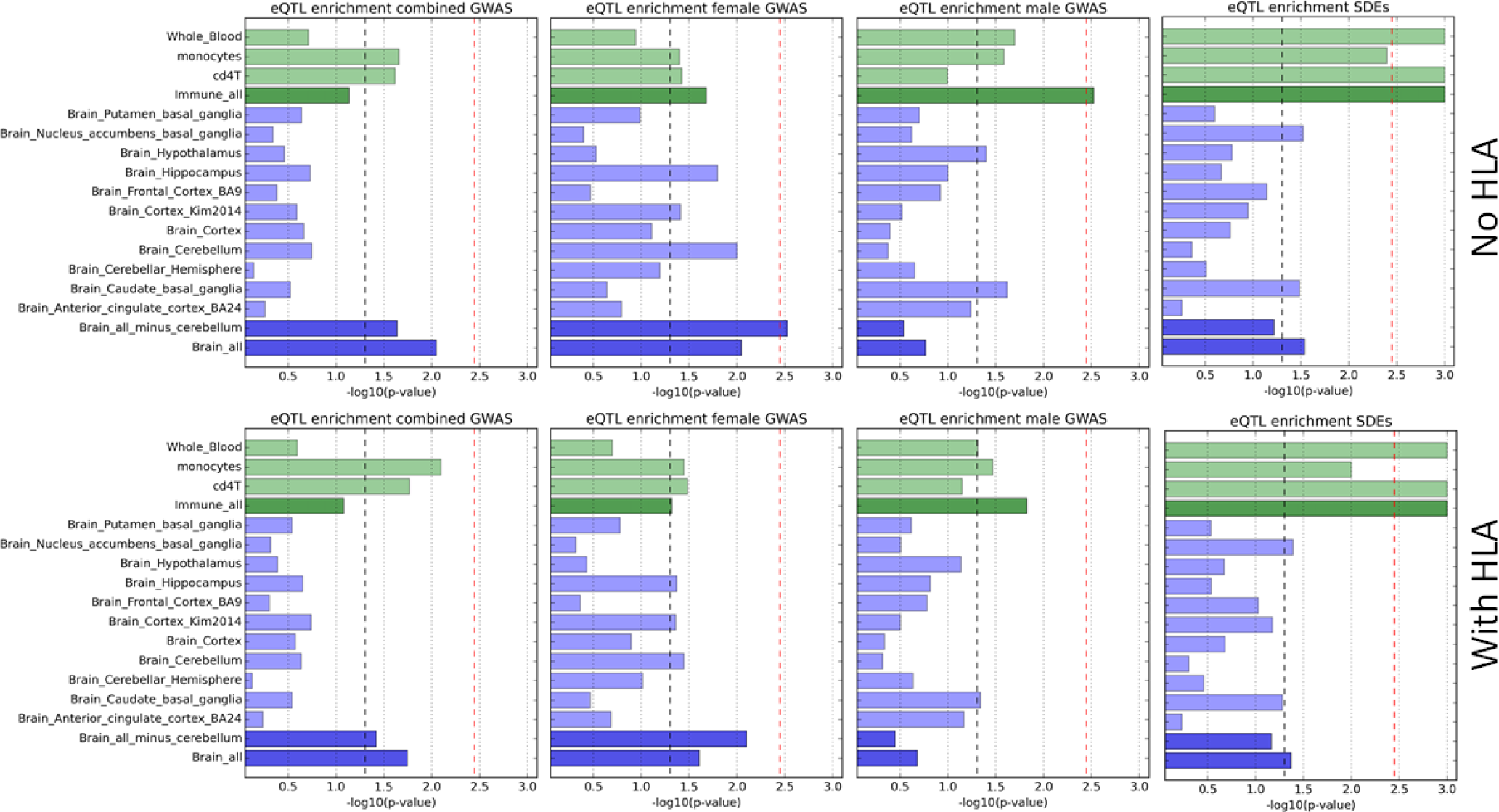
eQTL enrichment in the brain and immune tissues for combined, female-specific, male-specific top associations (10^−3^) and SNPs with Sexually Dimorphic Effect (SDEs), excluding and including SNPs in the HLA region. Only variants with more than 500 individuals in the GWAS are included here. Light green bars represent each immune tissue or cell type: whole blood, monocytes, and cd4+ T cells, while the dark green represents enrichment in a combination of the three immune tissues. Light blue bars represent each brain tissue, while the dark blue represents enrichment in a combination of ten brain tissues, or all ten brain tissues minus cerebellum. The black dashed line represents a p-value of 0.05. The red dashed line represents the significant p-value threshold (0.00357) after accounting for 14 eQTL datasets tested.

To characterize the function of the eGenes for those SNP sets showing a significant enrichment, we performed Gene Ontology (GO) enrichment analysis for biological processes and molecular functions using GSEA. Here we report the top two significant molecular processes and biological processes for the phenotypes showing significant enrichment of eGenes (full list in Supplementary Table 6). For SDEs, the immune eGenes showed the strongest enrichment for oxidoreductase activity (q=3.94e-04).) and hydrolase activity acting on ester bonds (q=3.94e-04) molecular functions, as well as organonitrogen compound biosynthetic process (q=7.77e-07) and cellular amide metabolic process (q=7.77e-07) biological processes. For top female associations, the brain eGenes showed enrichment only for two molecular function categories: ribonucleotide binding (q=1.38e-02) and cytoskeletal protein binding (q=3.06e-02). For top male associations, the immune eGenes showed an enrichment for structural constituent of ribosome (q=4.62e-02) molecular function, as well as organonitrogen compound metabolic process (q=2.66e-02). The reported q-values are the FDR adjusted significance value after correcting for multiple testing.

### Little overlap of OCD SDEs and anthropometric traits SDEs

Previous work has revealed enrichment of anthropometric traits SDEs (ASDEs) among top autism (ASD) and bipolar disorder (BIP) associated genetic variants, suggesting that the same mechanisms acting on secondary sex characteristic differences later in life may influence the risk of developing neurodevelopmental phenotypes such as ASD and BIP via pleiotropic effects (Mitra et al. 2016). There was little overlap and no significant enrichment for ASDEs among the clumped top combined, female-specific, male-specific GWAS associations, or OCD SDEs (Supplementary Figure 8).

### Males and females demonstrate similar levels of genetic correlation between OCD and other complex traits

As the lower bounds on the genetic correlation estimate of OCD between sexes ranged from 0.49-0.73, we explored whether males and females demonstrate differential genetic correlations between OCD and 30 traits (Supplementary Table 7) which may play a role in OCD development. Our analysis was limited by availability of combined male and female summary statistics only for the majority of the traits we tested in the correlation. The traits chosen for analysis included (1) neuropsychiatric phenotypes and behavioral traits (many of which exhibit sexually dimorphic characteristics), (2) traits which overlap with known sexually dimorphic clinical symptoms in OCD (e.g. smoking, eating disorders-anorexia, and body mass index), (3) brain structure volumes, and (4) reproductive behavior (age at first birth and number of children ever born).

We observed a significant genetic correlation of the combined male-female OCD sample with Tourette syndrome (r_g_=0.38, se=0.12, p=1.3e-03), anorexia (r_g_=0.59, se=0.15, p=7.85e-05), bipolar disorder (r_g_=0.56, se=0.11, p=6.10e-07), schizophrenia (r_g_=0.37, se=0.07, p=7.77e-07), neuroticism (r_g_=0.31, se=0.07, 3.38e-05), age at first birth (r_g_=0.37, se=0.07, 4.83e-07), and number of children ever born (r_g_=−0.35, se=0.09, p=6.66e-05). Several traits (bipolar disorder, schizophrenia, and neuroticism) exhibited a significant genetic correlation with female OCD, but not male OCD, possibly due to sample size (Supplementary Table 7).

In a sex-stratified genetic correlation analysis with neuropsychiatric and behavioral traits, none of the traits exhibited a statistically significant difference between female OCD x trait and male OCD x trait correlations (Supplementary Table 7), although Tourette syndrome, ADHD, anxiety, and bipolar disorder showed a trend towards a difference. We saw no statistically significant differences in the genetic correlation between sexes with body mass index, anorexia, smoking, brain structures volume or reproductive traits.

## Discussion

Obsessive-compulsive disorder is one of many neuropsychiatric traits exhibiting sexual dimorphism in both age of onset and presentation of symptoms. Overall, we find evidence for minor differences in the genetic architecture of OCD between the sexes, which suggests that heterogeneity from sex differences is not a significant contributor to loss of power in standard GWAS for OCD. Specifically, we report that the genetic correlation is high between males and females, and heritability estimates are not different between the sexes. Despite this, we observed differences in enrichment of functional annotations between variants with very different effects across the sexes. This finding suggests that although the study is still underpowered to detect these minor differences at the individual variant or gene level, we observe evidence of a portion of sexually dimorphic biology underlying risk for OCD. These results hold promise for discoveries in future studies with larger sample sizes. We expect the approaches developed here to enable a deeper understanding of how genetic variants may regulate biological processes influencing sex-biased phenotypes.

Male and female estimates of OCD heritability were nearly identical. This suggests that males and females share the same threshold of genetic liability for the development of OCD, consistent with previous reports meta-analyzing twin datasets (Polderman et al. 2015), indicating that the Carter Effect is not a major driver of sexual-dimorphism in OCD. We also identified a substantial genetic correlation for OCD risk between males and females. Furthermore, we observed a significant genetic correlation between OCD and several complex traits, including for several previously untested phenotypes, such as age at first birth and number of children born. While some traits (bipolar disorder, schizophrenia, and neuroticism) exhibited a genetic correlation with female OCD, we did not observe a significant correlation in males, likely due to sample size. Additionally, we observed no significant differences between male and female genetic correlations to any trait. We detected no genome-wide significant associations in either the sex-stratified GWAS or the genotype-sex interaction analyses. Partitioned heritability analysis indicates that the X chromosome contributes to the polygenic signal, but not more or less than expected given its size.

Despite the lack of significant associations in the sex-stratified GWAS or genotype-sex interaction analysis, we noted that the top GWAS associations were not the same between the sexes. Furthermore, we observed that SNPs with the greatest heterogeneity in effect size between males and females were enriched for gene regulatory function (eQTLs) in immune tissues, potentially implicating the immune system in sexual dimorphism of OCD. This finding is consistent with the known role of the immune system in several neuropsychiatric and neurodegenerative traits (Schizophrenia Working Group of the Psychiatric Genomics Consortium 2014; Marsh et al. 2016; Heneka, Golenbock, and Latz 2015; Furtado and Katzman 2015a, [b] 2015; Murphy, Sajid, and Goodman 2006). Recently, a small-scale whole-exome sequencing study showed that OCD families may have a higher rate of *de novo* nonsynonymous single-nucleotide variants in genes enriched for neurodevelopmental and immunological processes (Cappi et al. 2016). Alterations in the immune system and its function, including immune cell composition (Marazziti et al. 1999; Kawikova et al. 2007) and cytokine levels have been previously reported in individuals with OCD and are summarized by Murphy et al (Murphy, Sajid, and Goodman 2006). For example, a higher prevalence of OCD has been reported for patients with systemic lupus erythematosus (Slattery et al. 2004) and multiple sclerosis (Miguel et al. 1995). A high incidence of OCD and tics has also been reported in children who have had group A streptococcus infection and coined “pediatric autoimmune neuropsychiatric disorders associated with streptococcus: (PANDAS)” (Swedo et al. 1998; Murphy et al. 2012; Snider and Swedo 2004; Swedo et al. 2012). Symptom presentation in PANDAS is influenced by sex, with females more likely to present with chorea-like movements, and males more likely present with tics. Furthermore, males tend to present with tics, OCD and Sydenham's chorea at an earlier age (Swedo et al. 1998; Leonard et al. 1992; Carapetis and Currie 1999). Interestingly, in our study associations from the male GWAS showed a significant enrichment for eQTLs in immune tissues, and associations from the female GWAS were enriched for eQTLs in brain. Although a modest enrichment for eQTLs in immune tissues is also seen in female top associations, it is not significant after multiple testing correction. These findings suggest a potential difference in biology contributing to OCD in each sex.

Several limitations for this study should be noted, including sample size, and ascertainment strategies that may bias towards earlier age of onset which could result in uneven representation of disease subclasses among males and females. It has been reported that early-onset OCD is more heritable than adult-onset (Davis et al. 2013; Nestadt et al. 2000; van Grootheest et al. 2005), implicating genetic differences in early‐ and adult-onset OCD. Thus, uneven representation of males and females in the early‐ and adult-onset OCD groups could have led to measurement of equal heritabilities (i.e. in case of a comparison of early-onset males with adult-onset females, both of which should have a higher genetic burden for OCD). Although the majority of cases in this study are pediatric, there are adult-onset cases as well and a larger number of female cases may introduce a bias. Finally, the lack of detailed clinical data limits our ability to address many important questions related to symptom type, symptom severity, and age of onset. These limitations support the need for larger OCD datasets phenotyped in greater detail to delve deeper into examining the genetic contribution to sex differences of OCD.

Several recent studies (Rawlik, Canela-Xandri, and Tenesa 2016; Ge et al. 2017), have reported sexually dimorphic heritability, providing evidence for a sexually dimorphic liability threshold model for several human phenotypes. However, similar to other studies investigating sex-specific liability thresholds (Traglia et al. 2017; Martin et al. 2017; Mitra et al. 2016), we do not detect a difference in OCD genetic liability between the sexes. Sex-specific genome-wide significant associations have been identified in studies of ASD and ADHD, demonstrating the value of increasing sample size for the study of sexually dimorphic genetic effects.

## Supplementary Methods

### Sample and genotype quality control and imputation

To increase the power of the individual cohort GWAS, the alike sub-populations were combined i.e., OCGAS Europeans with IOCDF-GC Europeans, OCGAS Ashkenazi Jewish with IOCDF-GC Ashkenazi Jewish, and OCGAS Trios with IOCDF-GC Trios) after performing quality control for study-specific batch effects. Given that some of the trios were extracted from larger pedigrees, only one trio from each large family was retained. The trio data set included non-European individuals, which were removed prior to analysis. To identify study-specific batch effects, we performed principal component analysis to confirm no divergence of the two study populations on the first four principal components (PCs) (Supplementary Figure 2) and performed pseudo case-control GWAS by assigning case status to controls in one study (e.g. IOCDF-GC) and control status to controls in the other (OCGAS). There were no more significant associations than expected by chance when the SNPs with a missing genotype > 0.02 were removed. Those SNPs have a dosage that falls between 0.2 and 0.8 and thus cannot certainly be assigned to one genotype versus another, and thus may erroneously implicate a batch effect. Filtering for SNPs at various missingness threshold (from 0.01 to 0.1) did not have an effect of the GWAS result.

### Assessment of heterogeneity from sex-stratified GWAS

In addition to using the Z-score, we also estimated heterogeneity using Cochran's Q statistic, defined as the weighted sum of squared differences between individual study effects (in our case male and female variant effect sizes) and the pooled effect across studies, with the weights being those used in the pooling method, as implemented in METASOFT (Han and Eskin 2011). Because there was a high correlation between the two statistics, we utilized the Z-score in all subsequent analyses.

### Heritability estimates and genetic correlation

When assessing the contribution of the X chromosome to the heritability of a phenotype, it is important to consider the imbalance of the X chromosome dosage between males (with one copy of the X) and females (with two copies of the X). In sex-stratified analysis (female cases vs controls, and male cases vs controls) the dosage is balanced. However, when performing cross-sex comparison of heritability (i.e. males vs females) for X chromosome variants, a dosage compensation term is added to each pair of male:female co-variances to account for the dosage difference, which effectively models the X-linked genetic variance for females to the half that of males. Thus, for the cross-sex comparison analysis, the X chromosome genetic relatatedness matrix (GRM) was corrected by adding a dosage compensation flag in GCTA before running bivariate REML analysis.

## Supplementary Tables Legends

Supplementary Table 1. Sample size and the number of single nucleotide polymorphisms (SNPs) analyzed for each of the five (European, Ashkenazi Jewish, Dutch, South African, and Trios) datasets meta-analyzed. Number of cases and controls, and SNPs analyzed are shown for the sex-stratified and the combined sample.

Supplementary Table 2. Description of eQTL datasets used for eQTL enrichment analysis. Dataset source project, tissue type, sample size, reference and data url are shown.

Abbreviations: GTEx, Genotype-Tissue Expression project, ImmVar, Immune Variation project. Supplementary Table 3. Sex-stratified and combined heritability estimates for OCD from autosomes and the X chromosome utilizing various prevalence thresholds from 1-3% in 0.5% increments. Abbreviations: OCD, obsessive-compulsive disorder; LDSC, linkage disequilibrium score regression; GCTA, genome-wide complex trait analysis; N, number of individuals in the analysis; h^2^, SNP-heritability; SE, standard error.

Supplementary Table 4. Tables for eQTL enrichment in the brain and immune tissues for four groups: SDEs, combined OCD GWAS, male-specific and female-specific GWAS. Analysis was performed on variants with more than 500 individuals in the GWAS and an unfiltered set of variants (separate tab for each group and set of SNPs is shown). Within each tab, analysis of variants including and excluding the HLA region are shown. * denotes the condition with or without HLA where the result was statistically significant in one condition, but not the other. N_eGenes is the number of eGenes implicated by variants in each tissue tested, median and SD are the median number of eGenes and the standard deviation, respectively, in the 1000 matching sets. Enrichment p-value was calculated as the proportion of randomized sets in which the eGene count matches or exceeds the observed count in the list of the top male-specific, female-specific, combined GWAS variants or SDEs.

Supplementary Table 5. Tables showing eQTLs (rs id, chromosome, position, statistics associated with eQTL tests, associated eGene, and whether it is in the HLA region) among the top male-specific, female-specific, combined GWAS variants or SDEs. Each group (SDE, combined, male, and female) and eQTL dataset (GTEx, ImmVar, Cortex_kim2914) is listed in a separate tab.

Supplementary Table 6. Tables listing biological processes and molecular functions enriched among the eGenes implicated by top male-specific, female-specific, combined GWAS variants or SDEs in tissues showing significant eQTL enrichment (i.e. SDEs_immune, male_immune, and femle_brain). Gene set enrichment analysis tool (http://software.broadinstitute.org/gsea/index.jsp) was used to perform enrichment analysis, and the results are presented in each tab starting with “GSEA”.

Supplementary Table 7. Description of 30 GWAS studies used for genetic correlation analysis with OCD. Phenotype abbreviation, full name, reference, data source, sample size, sample prevalence, population prevalence, SNPs analyzed, phenotype statistics (heritability, LD score regression intercept, lambda GC), the sex-stratified and sex-combined genetic correlation statistics (r_g_, SE, p-value), as well as the male-female z-score are shown. Abbreviations: C, combined; M, male; F, female; r_g_, genetic correlation coefficient; se, standard error; SNPs, single nucleotide polymorphisms; NA, not applicable, OOB, out of bounds.

**Supplementary Figure 1.**
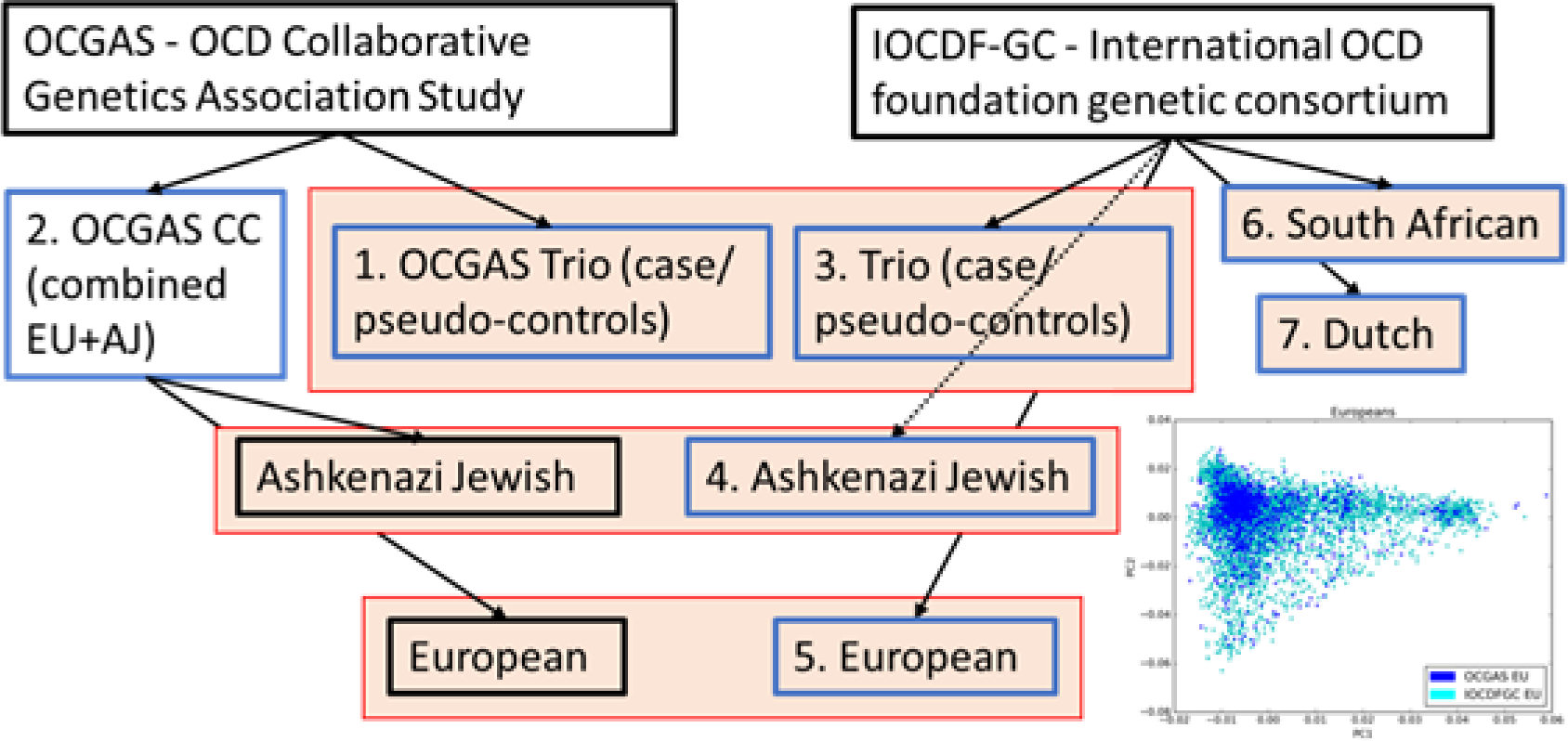
Details of study cohorts. The total cohort in this study is comprised of seven sub-populations collected as part of the OCD Collaborative Genetics Association Study (OCGAS) and the International OCD foundation genetic consortium (IOCDF-GC). The like populations from these two studies (trios, Ashkenazi Jewish, and European) were combined after extensive quality control to assess for study batch effect (Supplemental Methods). The inset figure of genetic principal component 1 (x-axis) vs principal component 2 (y-axis) for the European cohorts color coded by study (OCGAS:dark blue, IOCDF-GC:light blue) shows that the two studies do not form separate clusters, as would be indicative of a batch effect.

**Supplementary Figure 2.**
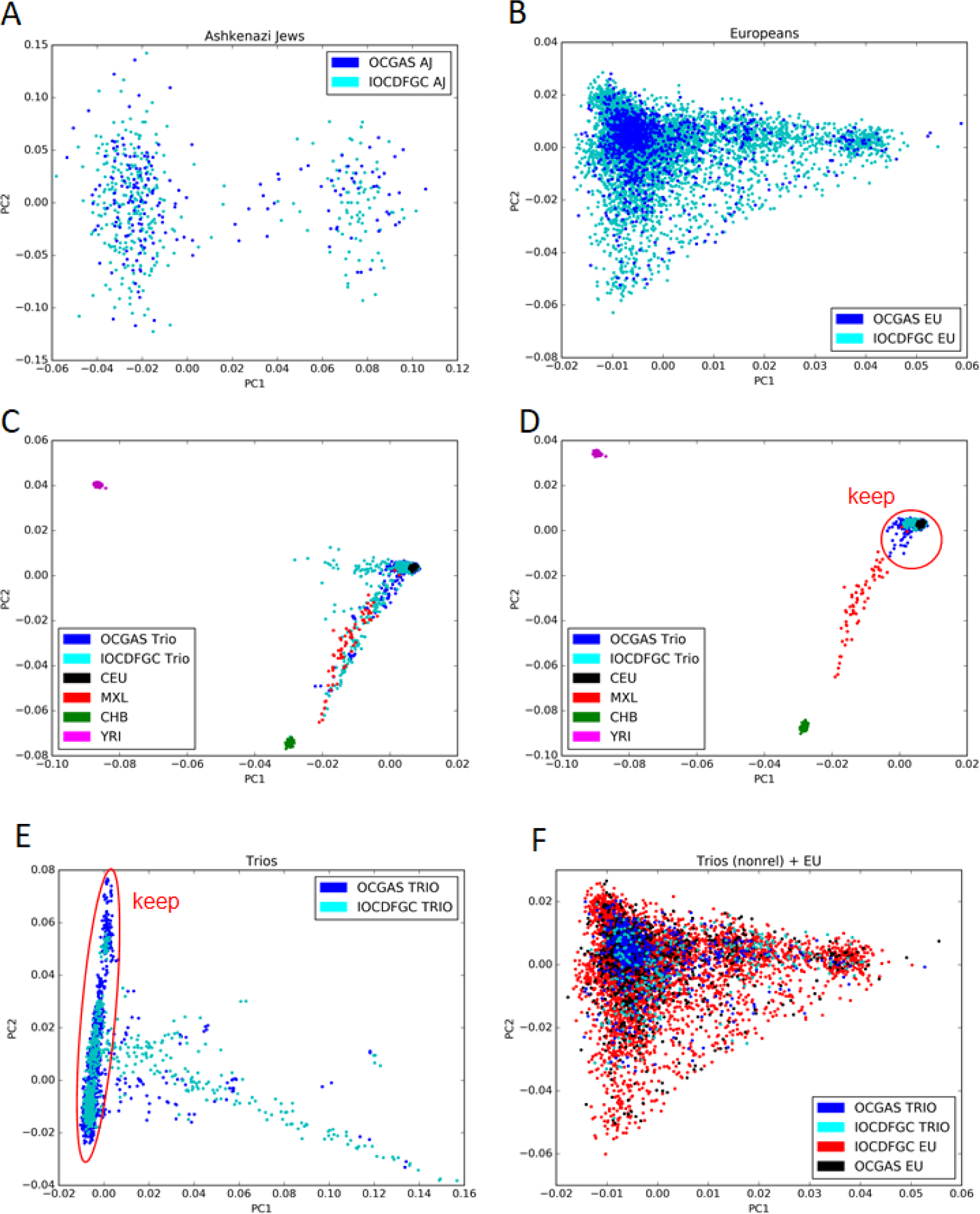
Principal component (PC) analysis plots for combined datasets. Genetic PC1 (x-axis) versus PC2 (y-axis) scatter plots indicate no batch effect when combining like cohorts: (A) Ashkenazi Jewish, (B) Europeans, and (F) Europeans with Trio cases. Plots C-E indicate how outliers were identified and removed from the trio subpopulation prior to combining with the European cohort (f) for heritability analysis. Abbreviations: OCGAS-OCD Collaborative Genetics Association Study; IOCDFGC-International OCD foundation genetic consortium; CEU-Utah Residents with Northern and Western European Ancestry; MXL-Mexican Ancestry from Los Angeles, USA; CHB-Han Chinese in Bejing, China; YRI-Yoruba in Ibadan, Nigeria.

**Supplementary Figure 3.**
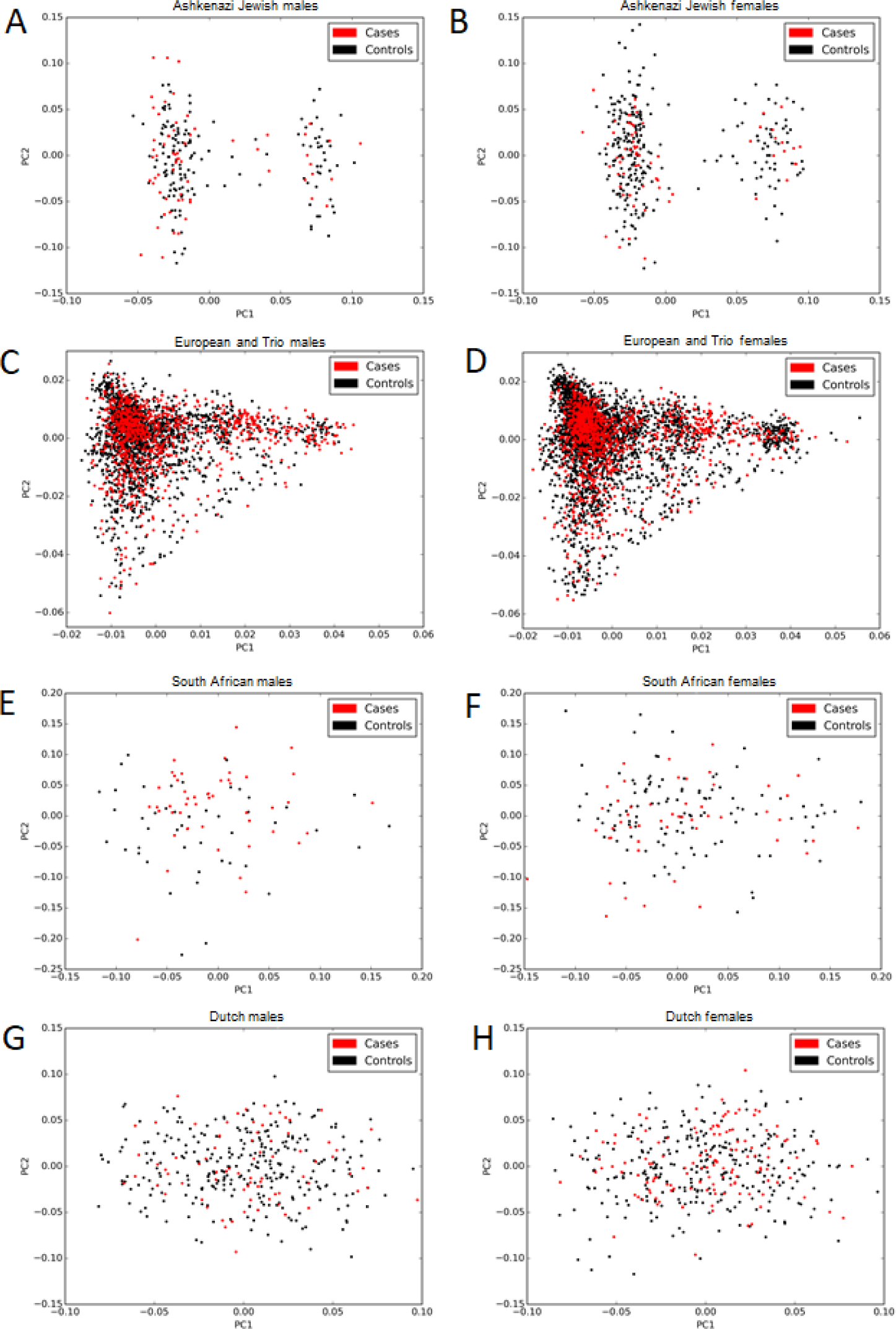
Principal component (PC) analysis plots stratified by sex. Genetic PC1 (x-axis) versus PC2 (y-axis) scatter plots stratified by sex indicate well-matched cases (red dots) and controls (black dots) for Ashkenazi Jewish (A-B), European and Trio (C-D), South African (E-F), and Dutch (G-H) sub-populations.

**Supplementary Figure 4.**
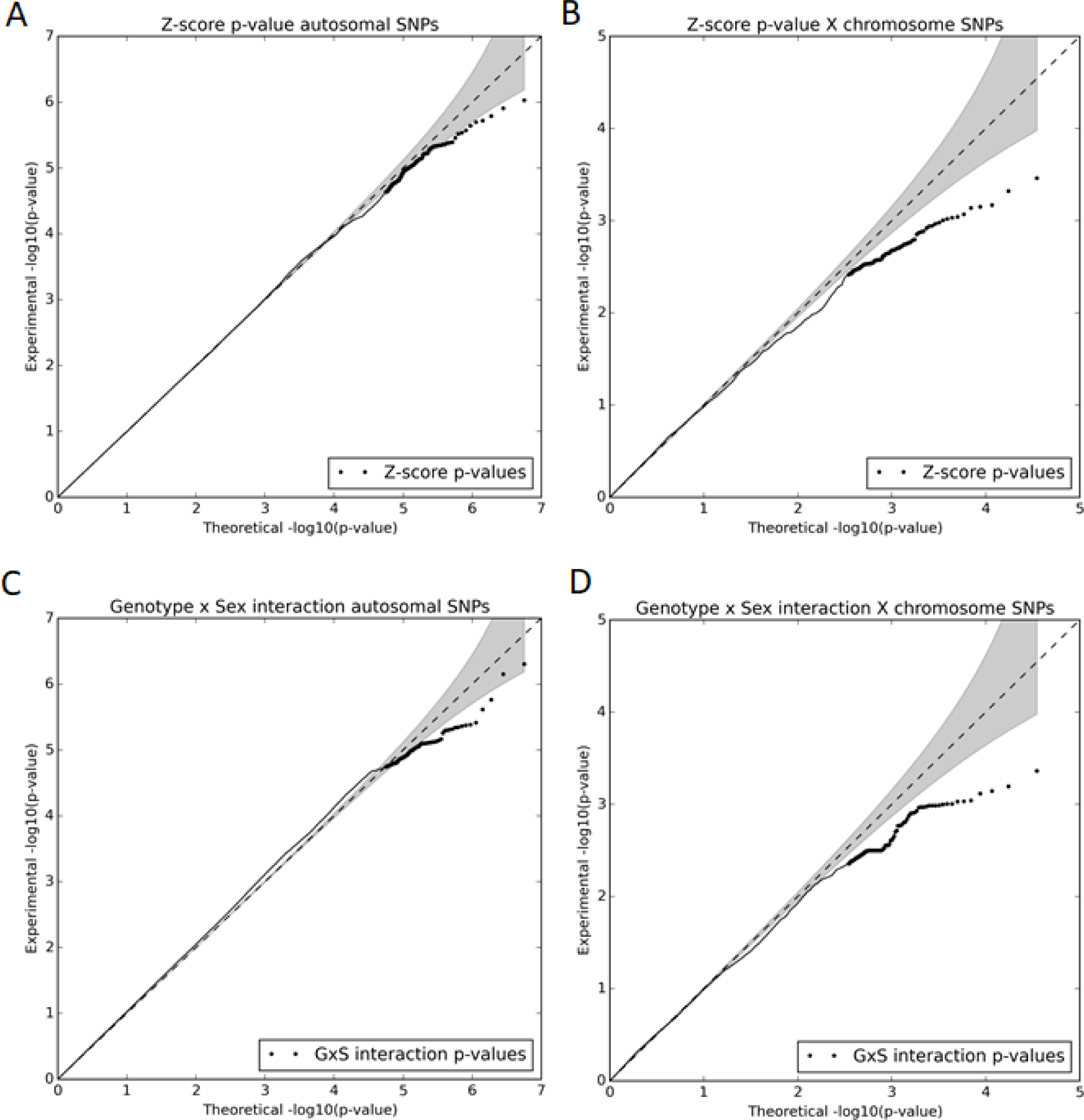
Quantile-quantile plots of the p-value distribution for Z-score, assessing difference in betas between the sexes, for the autosomal SNPs (A) and X chromosome SNPs (B), and the p-value distribution of the genotype-sex interaction term for the autosomal SNPs (C) and X chromosome SNPs (D). Only the variants with more than 500 individuals in the GWAS are included here.

**Supplementary Figure 5.**
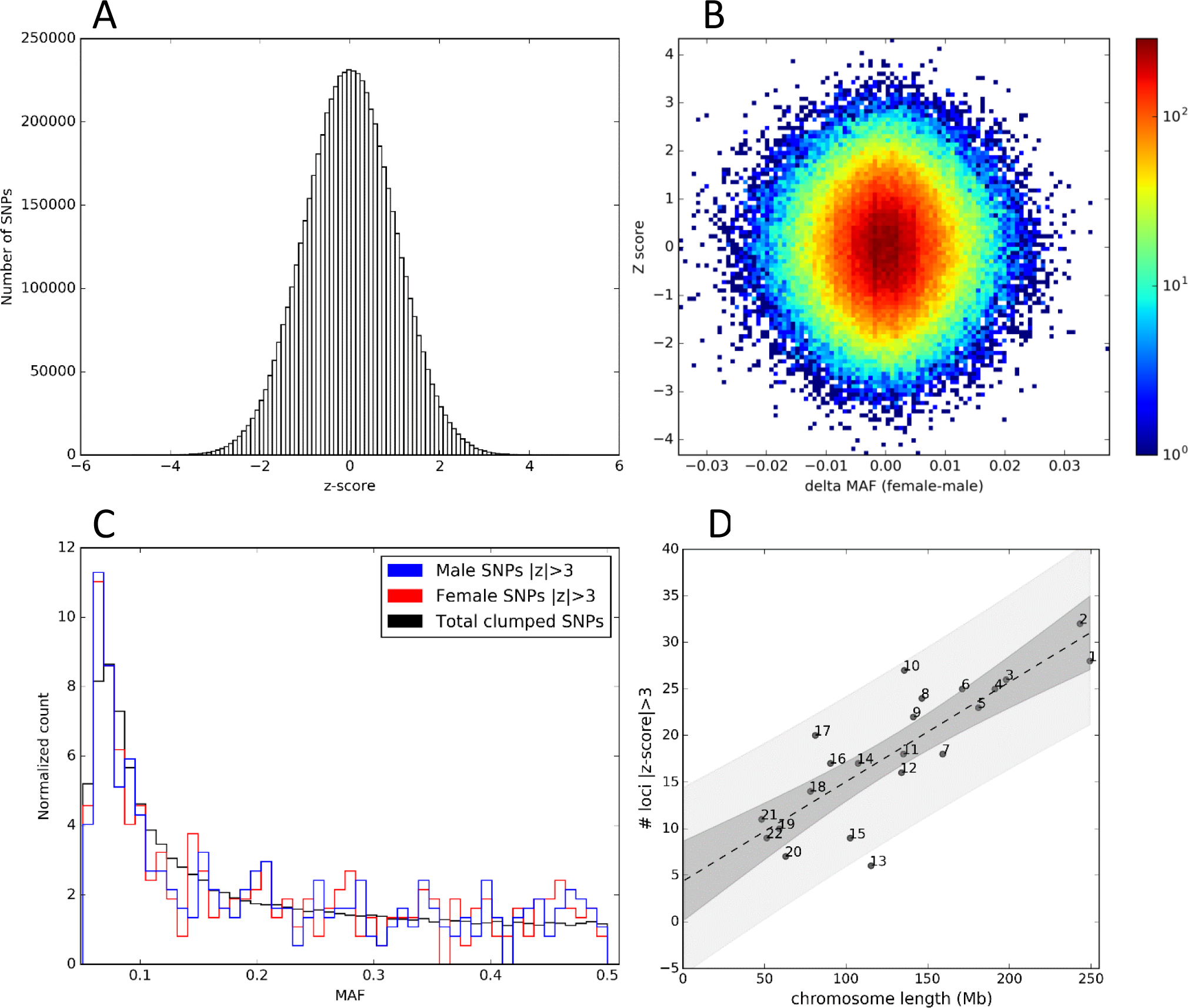
Characteristics SNPs with Sexually Differential Effect (SDEs). (A) Z-score distribution for all autosomal SNPs tested in the GWAS. (B) Scatter plot of Z-score versus difference in allele frequency between males and females for a set of clumped SNPs (n=167,450; r^2^=0.2, 500kb window). (C) MAF distribution by sex for SNPs with |Z-score| > 3 in the clumped set and for the all the SNPs in the clumped set (black). (D) There are autosomal 414 loci with |Z score| > 3 distributed across the genome proportional to chromosome length. Each dot is labeled with chromosome number. Dashed line represents the regression line (Pearson's r = 0.83, p=1.48e-06). The darker grey area represents the 95% confidence interval for the regression, while the lighter grey area represents the prediction interval, indicating that there is a 95% probability that the real value for the number of loci for a given chromosome length lies with the prediction interval. For B-D only the European cohort was used.

**Supplementary Figure 6.**
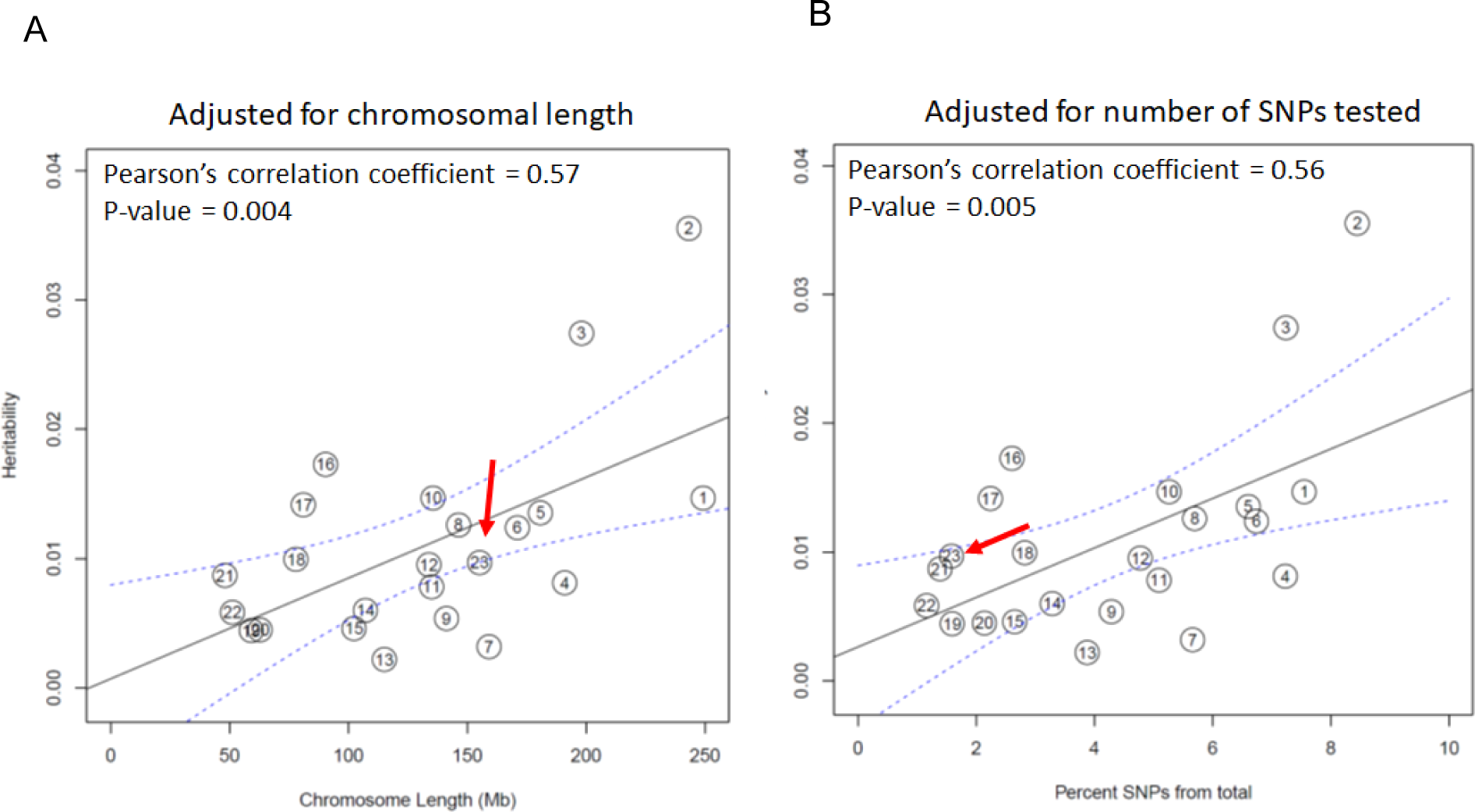
Partitioned OCD heritability by chromosome in the combined dataset. Heritability of OCD (y-axis) is plotted against chromosome length (A and percent of SNPs from total (B) on the x-axis. The black line represents heritability regressed on chromosome length (A) or percent of SNPs from total (B), with blue dashed lines represent the 95% confidence interval around the repression line. Red arrows point to chromosome 23 (X chromosome). The X chromosome contribution to heritability does not significantly deviate from expectation.

**Supplementary Figure 7.**
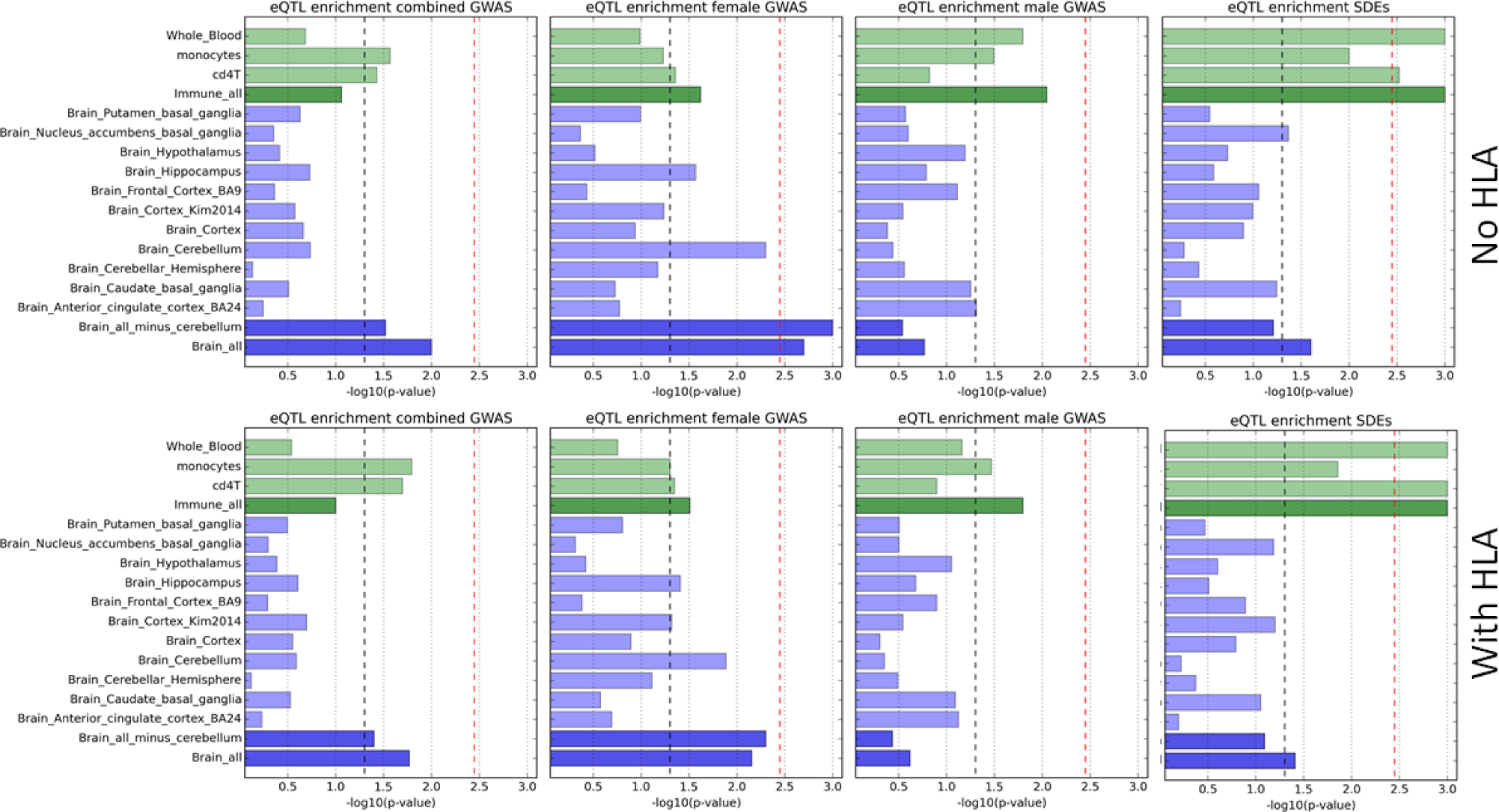
eQTL enrichment in the brain and immune tissues for combined, female-specific, male-specific top associations (10^−3^) and SDEs excluding and including SNPs in the HLA region for a set of SNPs without any restriction on the number of individuals contributing to the GWAS. Light green bars represent each immune tissue or cell type: whole blood, monocytes, and CD4+ T cells, while the dark green represents enrichment in a combination of the three immune tissues. Light blue bars represent each brain tissue, while the dark blue represents enrichment in a combination of ten brain tissues, or all ten brain tissues minus cerebellum. The black dashed line represents a p-value of 0.05. The red dashed line represents the significant p-value threshold (0.00357) after accounting for 14 eQTL datasets tested.

**Supplementary Figure 8.**
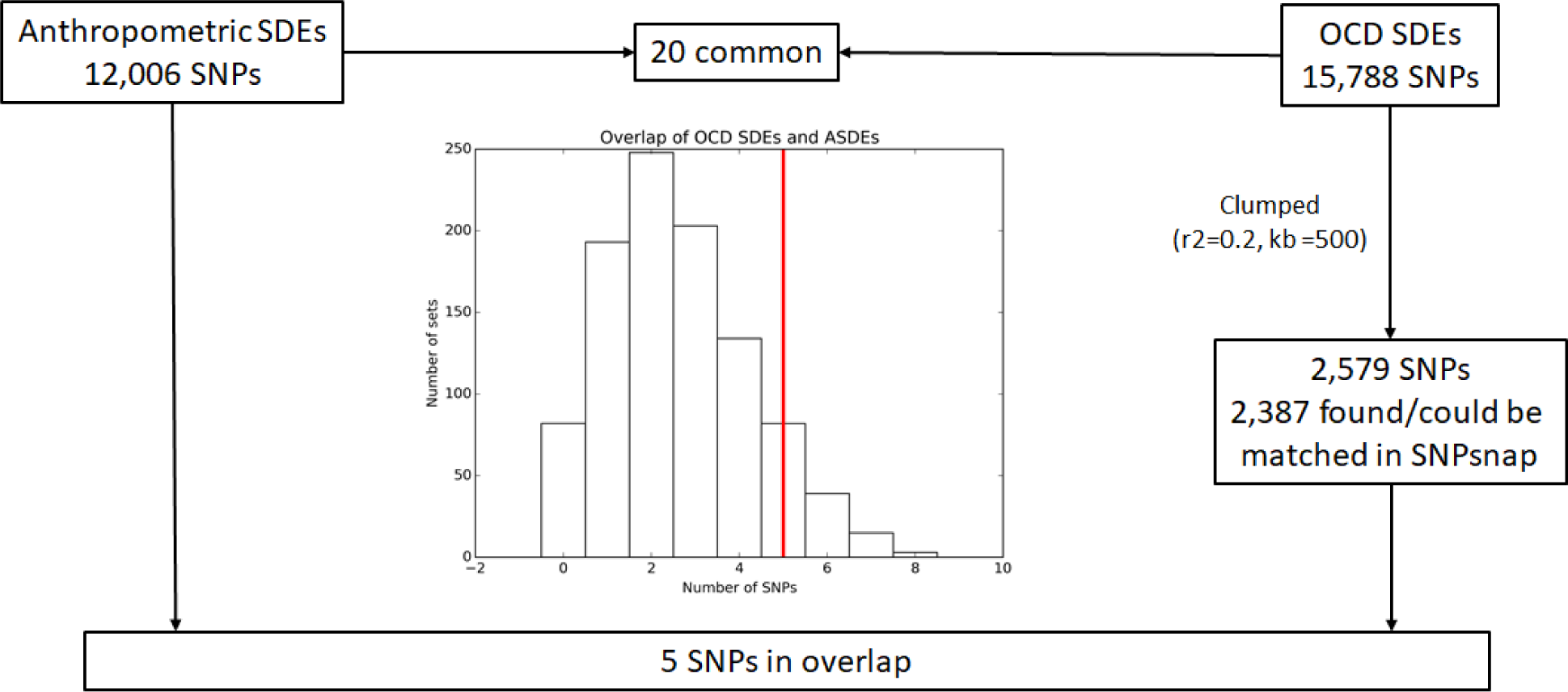
Schematic of enrichment of OCD SDEs among anthropometric traits SDEs (ASDEs). Twenty variants were common among ASDEs and OCD SDEs. When compared with 1000 matching sets for OCD SDEs, no enrichment was observed among ASDEs. The histogram shows the distribution of overlap of 1000 matching sets with SDEs, while the red bar represents the number of OCD SDEs after clumping that overlap ASDEs.

## Funding

The Conte Center for Computational Neuropsychiatric Genomics is funded by 3P50MH094267-04S1. The Center for Research Informatics is funded by the Biological Sciences Division at the University of Chicago with additional funding provided by the Institute for Translational Medicine, CTSA grant number UL1 TR000430 from the National Institutes of Health.

The OCD Collaborative Genetics Association Study (OCGAS) is a collaborative research study and was funded by the following NIMH Grant Numbers: MH071507 (GN), MH079489 (DAG), MH079487 (JM), MH079488 (AF), and MH079494 (JK). Yao Shugart and Wei Guo were also supported by the Intramural Research Program of the NIMH (MH002930-06)

The International Obsessive Compulsive Foundation Genetics Collaborative (IOCDF-GC) was supported by a grant from the David Judah Foundation (a private, non-industry related foundation established by a family affected by OCD), MH079489 (DLP), MH073250 (DLP), S40024 (JMS), MH 085057 (JMS), and MH087748 (CAM).

### Members of the IOCDF-GC and OCGAS consortia include (in alphabetical order)

Paul D. Arnold (IOCDF-GC)

Kathleen D. Askland (OCGAS & IOCDF-GC)

Cristina Barlassina (IOCDF-GC)

Laura Bellodi (IOCDF-GC)

OJ Bienvenu (OCGAS & IOCDF-GC)

Donald Black (IOCDF-GC)

Michael Bloch (IOCDF-GC)

Helena Brentani (IOCDF-GC)

Christie L Burton (IOCDF-GC)

Beatriz Camarena (IOCDF-GC)

Carolina Cappi (IOCDF-GC)

Danielle Cath (IOCDF-GC)

Maria Cavallini (IOCDF-GC)

David Conti (OCGAS)

Edwin Cook (IOCDF-GC)

Vladimir Coric (IOCDF-GC)

Bernadette A Cullen (OCGAS & IOCDF-GC)

Danielle Cusi (IOCDF-GC)

Lea K Davis (IOCDF-GC)

Richard Delorme (IOCDF-GC)

Damiaan Denys (IOCDF-GC)

Eske Derks (IOCDF-GC)

Valsamma Eapen (IOCDF-GC)

Christopher Edlund (IOCDF-GC)

Lauren Erdman (IOCDF-GC)

Peter Falkai (IOCDF-GC)

Martijn Figee (IOCDF-GC)

Abigail J Fyer (OCGAS & IOCDF-GC)

Daniel A Geller (IOCDF-GC and OCGAS)

Fernando S Goes (OCGAS)

Hans Grabe (IOCDF-GC)

Marcos A Grados (OCGAS & IOCDF-GC)

Benjamin D Greenberg (OCGAS & IOCDF-GC)

Edna Grünblatt (IOCDF-GC)

Wei Guo (OCGAS)

Gregory L Hanna (IOCDF-GC)

Sian Hemmings (IOCDF-GC)

Ana G Hounie (IOCDF-GC)

Michael Jenicke (IOCDF-GC)

Clare Keenan (IOCDF-GC)

James Kennedy (IOCDF-GC)

Ekaterina A Khramtsova (IOCDF-GC)

Anuar Konkashbaev (IOCDF-GC)

James A Knowles (IOCDF-GC and OCGAS)

Janice Krasnow (IOCDF-GC and OCGAS)

Cristophe Lange (OCGAS)

Nuria Lanzagorta (IOCDF-GC)

Marion Leboyer (IOCDF-GC)

Leonhard Lennertz (IOCDF-GC)

Bingbin Li (OCGAS)

K-Y Liang (OCGAS & IOCDF-GC)

Christine Lochner (IOCDF-GC)

Fabio Macciardi (IOCDF-GC)

Brion Maher (OCGAS)

Wolfgang Maier (IOCDF-GC)

Maurizio Marconi (IOCDF-GC)

Carol A Mathews (IOCDF-GC)

Manuel Matthesien (OCGAS)

James T McCracken (OCGAS & IOCDF-GC)

Nicole C McLaughlin (OCGAS & IOCDF-GC)

Euripedes C Miguel (IOCDF-GC)

Rainald Moessner (IOCDF-GC)

Dennis L Murphy (OCGAS & IOCDF-GC)

Benjamin Neale (IOCDF-GC)

Gerald Nestadt (OCGAS & IOCDF-GC)

Paul Nestadt (IOCDF-GC and OCGAS)

Humberto Nicolini (OCDF-GC)

Ericka Nurmi (OCGAS)

Lisa Osiecki (IOCDF-GC)

Carlos Pato (IOCDF-GC)

Michelle Pato (IOCDF-GC)

David L Pauls (IOCDF-GC and OCGAS)

John Piacentini (OCGAS & IOCDF-GC)

Danielle Posthuma (IOCDF-GC)

Ann E Pulver (OCGAS) H-D Qin (OCGAS)

Steven A Rasmussen (OCGAS & IOCDF-GC)

Scott Rauch (IOCDF-GC)

Margaret A. Richter (IOCDF-GC)

Mark A Riddle (OCGAS & IOCDF-GC)

Stephan Ripke (Psychiatric Genomics Consortium) Stephan Ruhrmann (IOCDF-GC)

Aline S. Sampaio (IOCDF-GC)

Jack F Samuels (OCGAS & IOCDF-GC)

Jeremiah M Scharf (IOCDF-GC)

Yin Yao Shugart (OCGAS & IOCDF-GC)

Jan Smit (IOCDF-GC)

Daniel Stein (IOCDF-GC)

S.Evelyn Stewart (IOCDF-GC and OCGAS)

Maurizio Turiel (IOCDF-GC)

Homero Vallada (IOCDF-GC)

Jeremy Veenstra-VanderWeele (IOCDF-GC)

Michael Wagner (IOCDF-GC)

Susanne Walitza (IOCDF-GC)

Y Wang (OCGAS & IOCDF-GC)

Jens Wendland (IOCDF-GC)

Nienke Vulink (IOCDF-GC)

Dongmei Yu (IOCDF-GC)

Gwyneth Zai (IOCF-GC)

## Computing resources

This work utilized the computational resources of the Center for Research Informatics' Gardner HPC cluster at the University of Chicago (http://cri.uchicago.edu) and the Dutch national e-infrastructure with the support of SURF Cooperative (https://userinfo.surfsara.nl/).

## Individuals

We thank Dr. Meritxell Oliva for assistance in preparation of GTEx eQTL data for analysis. We also wish to thank Drs. Carol Mathews, Jeremiah Scharf, and Edwin Cook for comments to improve the manuscript.

## Footnotes

URLs:

- Ricopili pipeline (Schizophrenia Working Group of the Psychiatric Genomics Consortium 2014) https://sites.google.com/a/broadinstitute.org/ricopili/
- Assocplots (Khramtsova and Stranger 2016) https://github.com/khramts/assocplots
- SHAPEIT (Delaneau, Zagury, and Marchini 2013) https://mathgen.stats.ox.ac.uk/genetics_software/shapeit/shapeit.html#home
- IMPUTE2 (Howie, Marchini, and Stephens 2011) https://mathgen.stats.ox.ac.uk/impute/impute_v2.html
- PLINK2 (Purcell et al. 2007) https://www.cog-genomics.org/plink2
- EIGENSOFT-smartpca (Price et al. 2006) https://data.broadinstitute.org/alkesgroup/EIGENSOFT
- METAL (Willer, Li, and Abecasis 2010) http://csg.sph.umich.edu//abecasis/Metal/
- METASOFT(Han and Eskin 2011) http://genetics.cs.ucla.edu/meta/
- GCTA (Yang et al. 2011) http://cnsgenomics.com/software/gcta/
- GSEA (Mootha et al. 2003; Subramanian et al. 2005) http://software.broadinstitute.org/gsea/index.jsp
- LDSC (B. K. Bulik-Sullivan et al. 2015; B. Bulik-Sullivan et al. 2015) https://github.com/bulik/ldsc
- SNPsnap (Pers, Timshel, and Hirschhorn 2015) https://data.broadinstitute.org/mpg/snpsnap/

